# Allele-Specific Quantification of Structural Variations in Cancer Genomes

**DOI:** 10.1101/048207

**Authors:** Yang Li, Shiguo Zhou, David C. Schwartz, Jian Ma

## Abstract

One of the hallmarks of cancer genome is aneuploidy, resulting in abnormal copy numbers of alleles. Structural variations (SVs) can further modify the aneuploid cancer genomes into a mixture of rearranged genomic segments with extensive range of somatic copy number alterations (CNAs). Indeed, aneuploid cancer genomes have significantly higher rate of CNAs and SVs. However, although methods have been developed to identify SVs and allele-specific copy number of genome (ASCNG) separately, no existing algorithm can simultaneously analyze SVs and ASCNG. Such integrated approach is particularly important to fully understand the complexity of cancer genomes. Here we introduce a new algorithm called Weaver to provide allele-specific quantification of SVs and CNAs in aneuploid cancer genomes. Weaver uses a probabilistic graphical model by utilizing cancer whole genome sequencing data to simultaneously estimate the digital copy number and inter-connectivity of SVs. Our simulation evaluation, comparison with single-molecule Optical Mapping analysis, and real data applications (including MCF-7, HeLa, and TCGA whole genome sequencing samples) demonstrated that Weaver is highly accurate and can greatly refine the analysis of complex cancer genome structure.

## 1 Introduction

Genome aneuploidy, in which abnormal copy numbers of alleles are present, is a hallmark of cancer [1, 2]. A large proportion of tumors are aneuploid and have undergone either arm-level somatic copy number alterations (CNAs) or even whole-genome duplications (WGD) [1, 3, 4]. In some types of cancer such as bladder cancer, ovarian cancer, and lung cancer, more than 50% of the tumors have undergone WGD [4]. Structural variations (SVs), including deletions, insertions, duplications, and rearrangements, can further modify the aneuploid cancer genome into a mixture of rearranged genomic segments with extensive range of CNAs. Indeed, aneuploid cancer genomes have significantly higher rate of CNAs as well as SVs [4]. A comprehensive and precise characterization of these changes is critical in understanding the evolution of cancer genome [5] and in interpreting cancer-specific gene expression and epigenetic alterations using high-throughput next-generation sequencing (NGS) data [6].

Allele-specific copy number of genome (ASCNG) analysis has been performed for SNP array data [13, 14] and recently for NGS data as well [9–12]. Separately, SV identification methods have also been developed for NGS data, such as [5, 6, 9, 11]. It is essential to ask how SVs interact with ASCNG and how different SVs interact with each other. The answers to such questions can help unravel the detailed cancer genome structure and its evolutionary history. However, integrative method specifically for simultaneously analyzing SVs and ASCNG has not been developed. Indeed, except arm-level gain/loss, the majority of somatic CNAs are associated with SVs [17]. It has been reported that analyzing CNAs around SV breakpoints can reveal the mutational forces causing particular cancer subtype [17–19]. Moreover, the integrated approach can further assist the variants phasing in different scales (both SNPs and SVs) in the context of complex cancer genome architecture.

In this paper, we introduce a novel computational method Weaver to identify allele-specific copy number of SVs (ASCNS) as well as the inter-connectivity of them in aneuploid cancer genomes. To our knowledge, this is the first method that can simultaneously analyze SVs and ASCNG. Under the same method framework, Weaver also provides base-pair resolution ASCNG. Note that in this paper we specifically focus on the quantification of SV copy numbers, which is also the key novelty of our method. Our framework is flexible to allow users to choose their own variant calling (including SV) tools. We use the variant calling results to build cancer genome graph, which is subsequently converted to a pair-wise Markov Random Field (MRF). In the MRF, the ASCNS and SV phasing configuration, together with genomic ASCNG, are hidden states in nodes and the observations contain all sequencing information, including coverage, read linkage between SNPs as well as between SV and SNPs. Therefore, our goal of finding the ASCNS and SV phasing together with ASCNG is formulated as searching the *maximum a posteriori* (MAP) solution for MRF. We apply Loopy Belief Propagation (LBP) framework to solve the problem.

Being an integrative graphical model to analyze SVs and CNAs that fits well with the complex nature of cancer genomes with reorganized chromosomes, Weaver’s novel contribution is trifold: (i) The method provides a quantitative measurement of SVs in cancer genome; (ii) It estimates the phasing/linkage information of different SVs using NGS data; (iii) The method generates the highly accurate base-pair resolution (note that all previous methods can only provide rough estimate of CNA boundaries, as in Appendix Fig. 1) ASCNG profiling in aneuploid cancer genomes, by simultaneously achieving (i) and (ii). Our simulation evaluation, comparison with single-molecule Optical Mapping analysis, and real data applications (on MCF-7, HeLa, and TCGA whole-genome sequencing datasets) demonstrated that Weaver is highly accurate and can significantly refine the analysis of complex cancer genomes.

**Figure 1:**
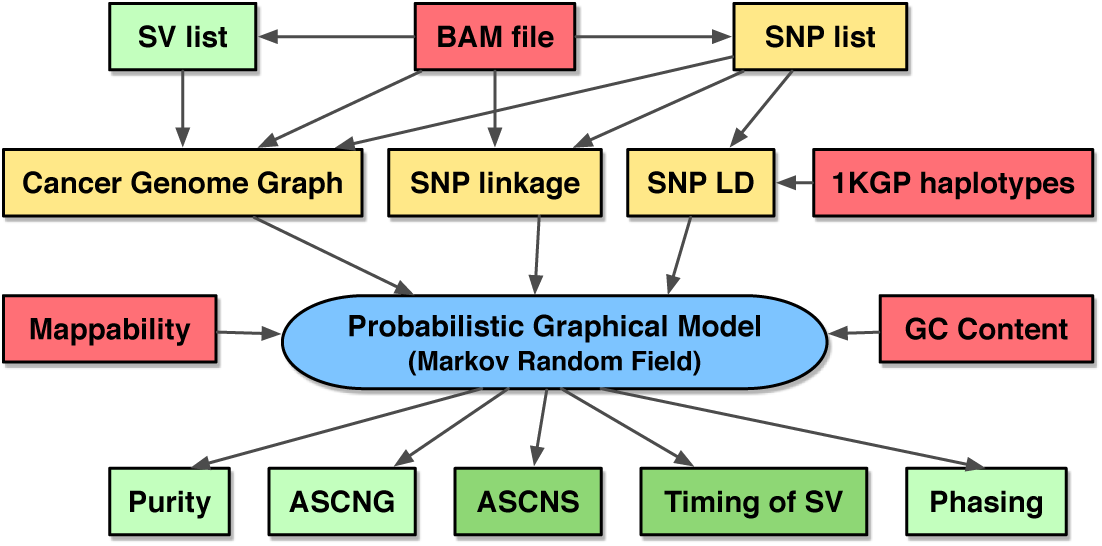
The method overview of Weaver. Red boxes represent input and green boxes represent output. Yellow boxes show intermediate results. Dark green boxes show the primary goals of Weaver that are novel and have never been tackled by other methods, while light green ones show ‘by-products’ of Weaver that have also been demonstrated to perform better in Weaver as compared to existing methods for these outputs.

## 2 Methods

The overview of the Weaver algorithm is shown in Fig. 1. The input of Weaver is the BAM file of aligned and unaligned reads from a particular tumor sample. If there is matched normal sample available, it will also be used (details in Section 2.3). The first step is to call variants (including both SNPs and SVs) based on the BAM file. Users can choose their own variant calling methods. The detailed description for preparing Weaver input is in the Appendix. In the Methods section here, we focus on the most important aspects of our work.

Using the intermediate results (yellow boxes in Fig. 1) including the cancer genome graph construction (Section 2.1), the Weaver MRF model will be built. By solving the MRF MAP function (Equation 4), Weaver generates output as shown in the green boxes in Fig. 1 (see Fig. 2E for example). Weaver source code is freely available and can be downloaded from: https://github.com/ma-compbio/Weaver.

**Figure 2:**
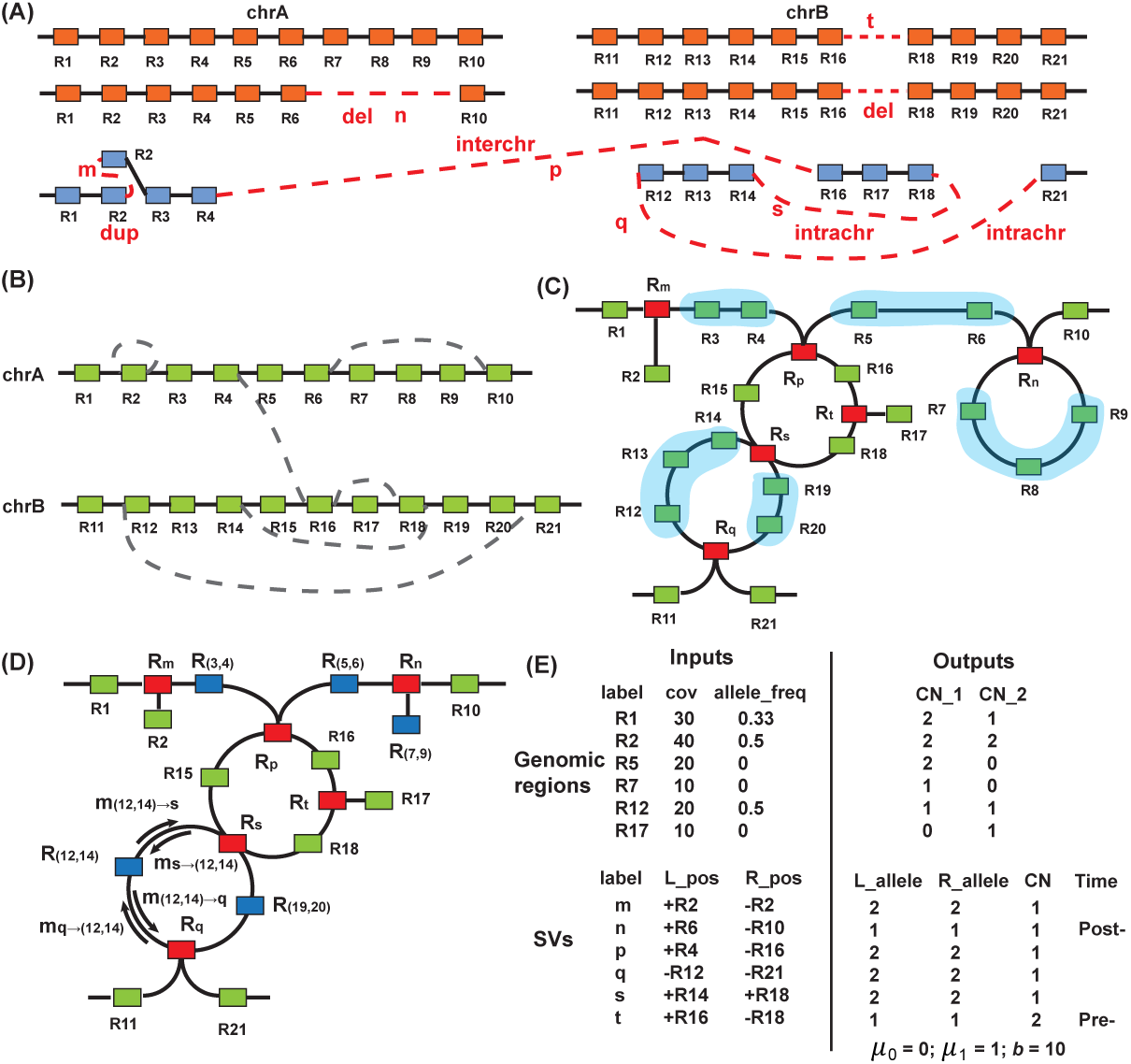
MRF model details. **(A)** Hypothetical cancer chromosomes with rearrangement structure hidden. Orange and blue segments represent paternal/maternal allele. Red dashed line represent linkages by SVs. **(B)** The cancer genome graph, constructed from (A), with nodes (boxes) representing genomic regions and edges representing reference (solid lines) or cancer (dashed lines) adjacencies. **(C)** MRF representation in Weaver. Red boxes represent *cancer nodes*(𝓡_*c*_) that have included SVs information; green boxes are the same with (B) and representing *genome nodes*(𝓡); the lines between *genome nodes* are *genome edges*(𝓔^*r*^); the lines between *cancer nodes* and *genome nodes* are *cancer edges*(𝓔^*c*^). **(D)** Blue boxes represent supernodes by clustering blue shaded chains of *genome nodes* as shown in (C). **(E)** Input and output of MRF are separated into genomic regions and SVs. For region *R*_1_, the input is observed coverage 30 and allele frequency 0.33; the output is 2 copies on allele 1 and 1 copy on allele 2. n is a post-aneuploid deletion with 1 copy and both breakpoints are on allele 1 of chrA. *t* is a pre-aneuploid deletion with 2 copies and both breakpoints are on allele 1 of chrB. SV *m*, *p*, *q* and *s* are from the allele that has not been duplicated.

### 2.1 Genome partitioning and cancer genome graph construction

We first select a default size *W* (e.g., 5kb) and partition the genome into non-overlapping regions as follows: (i) Breakpoints in the input SV set 𝓒 must be on region boundaries; (ii) Each region may contain no more than one SNP; (iii) The size of each region must be ≤ *W*. The number of regions from initial segmentation in Weaver ranges from 1.7 million to 2 million based on various datasets in this work, depending on the size of loss of heterozygosity (LOH) regions and the number of SVs. This is a combined strategy that utilizes both fixed window size and SV boundaries for segmentation. Since SV breakpoints and CNV boundaries do not always match, our proposed MRF models this probabilistically. Overall, this approach has the advantage to provide base-level ASCNG boundaries as compared to existing genome segmentation methods in copy number analysis, which typically use fixed segmentation size.

Given the segmentation of the genome and SV set 𝓒, we then build *cancer genome graph* 𝓖:= {𝓡, 𝓔} (Fig. 2B), with nodes representing genomic region sets (𝓡) and edges representing *reference adjacencies* (𝓔^*r*^) (solid lines in the figure) if two nodes are adjacent in the normal genome and *cancer adjacencies* (𝓔^*c*^) (dashed lines in the figure) if two nodes are adjacent in the cancer genome by SV *c* linkage. Edge configurations 𝓔 between node *R*_*i*_ and *R*_*j*_ can be represented as: (*δ*_*i*_*R*_*i*_ ~ *δ*_*j*_*R*_*j*_), *δ* ∈ {+, –}, with + and – representing the tail (right) and head (left) of a given genomic region *R*, e.g., (+*R*_*i*_ ~ –*R*_*i*+1_) ∈ 𝓔^*r*^, if *R*_*i*_ and *R*_*i*+1_ are adjacent regions from the same chromosome in the normal genome.

We then convert the original cancer genome graph 𝓖:= {𝓡, 𝓔} into Markov Random Field (MRF, 𝓜:= {𝓡, 𝓡_*c*_, 𝓔^*r*^, 𝓔^*c*^}), which is a widely used probabilistic graphical model to estimate joint probabilities. The MRF can be viewed as an undirected graph and the aggregated inference problem in Weaver given sequencing data can be viewed as a *maximum a posteriori* (MAP) problem with hidden states and observations explained in the following sections. Unlike conventional methods for estimating copy number changes based on hidden Markov models (HMMs), which are designed for sequential data and only consider the dependencies between ‘local’ variables, MAP solution of MRF model provides the most probable configuration of aneuploid cancer genomes with complex SVs, involving ‘global’ variable dependencies defined by long-range SVs (i.e., distal connections of variants). This is the main rationale of using MRF for our problem. In the following sections, we describe hidden states, observations, and formal function of the MRF MAP problem. Details on potential functions on nodes and edges are provided in the Appendix.

### 2.2 Hidden states 𝓗

For *i*^*th*^ *genome node R*_*i*_ ∈ 𝓡 ⊂ 𝓜, the hidden states are 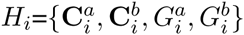, where 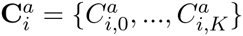 and 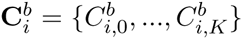 are vectors of non-negative integer numbers representing copy numbers for allele *a* and *b* of *k*^*th*^ population on *R*_*i*_, respectively. When *k* = 0, it stands for the fraction of normal cells. Note that although the Weaver algorithm is generic and in principle can be applied for multiple subclones (*K* > 1), in our current implementation, Weaver only processes tumor samples without significant subclonal structure (i.e., *K* = 1). We leave the cases for *K* > 1 with tumors with complex subclonal structure as future work.

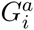 and 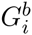 represent the genotype of allele *a* and *b* of *R*_*i*_, which is independent from subclone structure since only germline SNPs are considered. For convenience, we also set variable *C*_*i*, *k*_ as the overall copy number of *k*^th^ population on 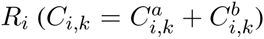. In our analysis of cancer genomes, which typically have highly amplified regions, we do not have a limit for *C*_*i*, *k*_, as done by previous CNA methods. The hidden copy number is bounded by the observation of sequencing depth on each region. Note that for regions with low mappability or extreme GC content, it is not reliable to infer hidden state space with observed local sequencing coverage; instead, we search the closest region and inherit its hidden state space setting, assuming that there is no dramatic state change between them. The hidden states on *cancer nodes* 𝓡_*c*_ are discussed in Appendix 5.4.

### 2.3 Observations 𝓞

For observation on 𝓡 ⊂ 𝓜, on *i*^th^ genomic region *R*_*i*_ ∈ 𝓡, the observation from the hidden state is the raw read coverage *O*_*i*_ on entire *R*_*i*_, which can be estimated by tools such as BEDTools [20] based on BAM file. For tumor sample with matched normal genome sequenced, we calculate 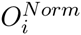 for the same *R*_*j*_, and normalize the *O*_*i*_ using: 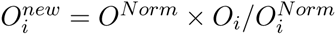, where *O*^*Norm*^ is the median coverage of the entire normal genome.

If *R*_*i*_ has SNP, 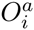 and 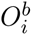 are the number of reads containing the SNP based on *a*/*b* allele, respectively, which can be obtained from SNP calling pipelines such as [21]. In practice, neither sequencing nor mapping is uniform across the genome. Here we consider two widely used factors, the GC-content and short read mappability (from UCSC). Using two HapMap samples NA18507 and NA12878, we split the human genome into consecutive 100bp bins and calculated the average mapping coverage on each bin. Among the bins that have unexpected low or high coverage as compared to the rest of the genome, more than 91% have either mappability < 0.6 or GC-content < 0.2 or > 0.6. Therefore, we label all *R*_*i*_ as not read-depth informative, if mappability < 0.6 or GC-content < 0.2 or > 0.6. The read depth of those uninformative regions are inherited from neighboring regions.

Regarding observation on 𝓔^*r*^ ⊂ 𝓜, within two adjacent genomic regions *R*_*i*_, *R*_*i*+1_ ∈ 𝓡, there are two independent observations for their genotype linkage.

(i) We assume the genotypes on *i* and *i* + 1 are 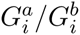 and 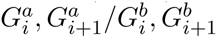, respectively. We define the Linkage Disequilibrium (LD) score for the phasing configuration 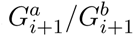 as:

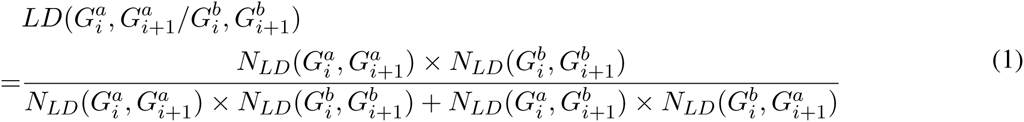

where 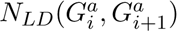 is the number of phased haplotypes (total number 1092 × 2 in phase 1) in 1KGP with genotype 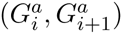. Other genotype configurations can be similarly calculated.

(ii) Similarly, we define the read linkage score for the phasing 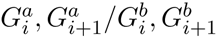 as:

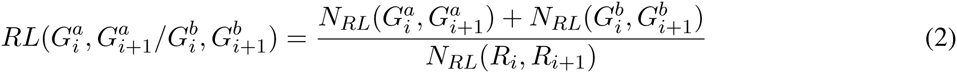

where *N*_*RL*_(*R*_*i*_, *R*_*i*+1_) is the total number of reads covering genomic regions (*R*_*i*_, *R*_*i*+1_) and 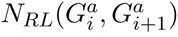 is total number of reads covering 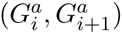. If there are no reads covering (*R*_*i*_, *R*_*i*+1_) (*N*_*RL*_(*i*, *i* + 1) = 0), *RL* = 0.

Therefore, we define genotype linkage as

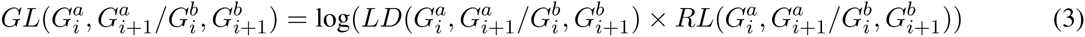

In real data application, we have found that *RL* and *LD* correlate very well. For example, in the MCF-7 genome analysis (see Results section), when we chose SNP pairs with 100% *RL* support as gold standard, we found AUC= 0.9964 using *LD* scores.

### 2.4 Markov Random Field model 𝓜

After we convert 𝓖 into MRF 𝓜 using steps in Appendix 5.6, the MRF MAP problem is given by:

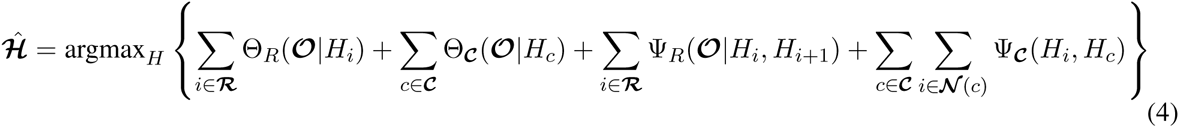

where Θ_*R*_(𝓞|*H*_*i*_) is the *genome node* (green box in Fig. 2C, Appendix 5.2) potential function. Θ_𝓒_(𝓞|*H*_*c*_) denotes constraint function in *cancer nodes* (red box in Fig. 2C, Appendix 5.4). Ψ_*R*_(𝓞|*H*_*i*_, *H*_*i*+1_) is the *genome edge* (link between green boxes in Fig. 2C, Appendix 5.3) function, providing pairwise constraints of hidden states of neighboring nodes *R*_*i*_, *R*_*i*+1_. Ψ_𝓒_(*H*_*i*_, *H*_*c*_) is the *cancer edge* (link between green and red box in Fig. 2C, Appendix 5.5) potential function. 𝓝(*c*) stands for the index of *genome nodes* linked to SV *c*.

The general MRF MAP problem is computationally intractable [22]. Several approximation approaches have been proposed to solve this problem. Here we utilize Belief Propagation (BP) to solve the MRF MAP problem. BP was originally proposed for graphs without cycle [23], in which case the fixed point of max-product belief propagation algorithm are also the assignment of MAP [24]. When applying on graph with arbitrary topology, the Loopy Belief Propagation (LBP) can still approximate well to the MAP configuration [25]. With 𝓗̂ estimated, the 𝓗̂_*i*_ on *genome node* provides base-pair resolution ASCNG and 𝓗̂_*c*_ on *cancer node* provides estimation of ASCNS.

We use LBP to find the MAP configuration of MRF [25]. The message updating rule from node *R*_*j*_ to node *R*_*i*_ (as illustrated in Fig. 2D) at (*t* + 1)^*th*^ iteration is:

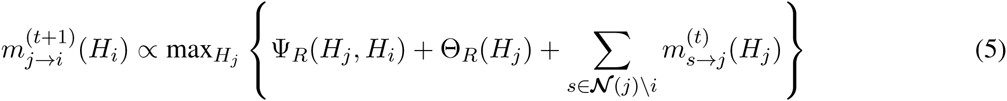

where 𝓝(*j*)\*i* stands for index of all the nodes linked to node *j*, except for node *i*. Note that the max-product form of message passing is used to get state configuration with MAP. The above function assumes *R*_*i*_ and *R*_*j*_ are *genome nodes*, corresponding potential will be replaced if *R*_*j*_, or *R*_*j*_ is *cancer node*.

The belief vector (max-marginal) is computed for each node at *t*^*th*^ iteration:

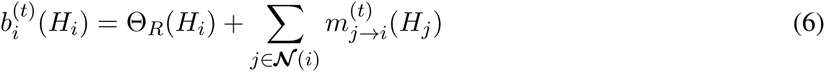

If convergence 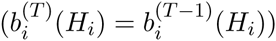 or the maximum iteration number is reached at *T*^*th*^ iteration, the final belief vector for each node is 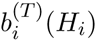. The set of 𝓗̂ that provides the largest belief: 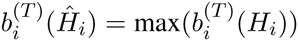 will be the MAP result for our problem. Since the message passing in BP is proportional to the number of nodes, we reduce the number of nodes using a procedure described in Appendix 5.7 in order to make the overall computation more efficient.

As illustrated in Fig. 2E, the final output of Weaver has three major parts: (i) the purity (*μ*_0_ and *μ*_1_) and haplotype level coverage *b*; (ii) ASCNG; (iii) ASCNS, as well as the timing of SVs with respect to chromosome amplification or deletion (aneuploid).

## 3 Results

### 3.1 Evaluation by simulation

We first evaluated the performance of Weaver on simulated datasets. The detailed steps on how the simulated data were generated are described in Appendix 5.8.

#### 3.3.1 Accuracy of estimating ASCNS in Weaver

Overall, Weaver identified 97.1% SVs with correct copy number and 95.7% of SVs were phased to correct allele. The timing of SV can be inferred with pre‐ and post-aneuploid SVs. We have correctly detected 97.3% pre‐ and 98.7% post-aneuploid SVs. SN and SP of reporting SV with specific copy numbers are summarized in Fig. 3A-B. The dispersion parameter *ϕ* is approximated by adding various degrees of random noises on original coverage from simulation data. With increasing noise level (larger dispersion *ϕ*), both SN and SP drop. However, based on our observation on the real datasets, the dispersion is typically a bit greater than 1, suggesting that Weaver should perform well on real cancer genome data.

**Figure 3:**
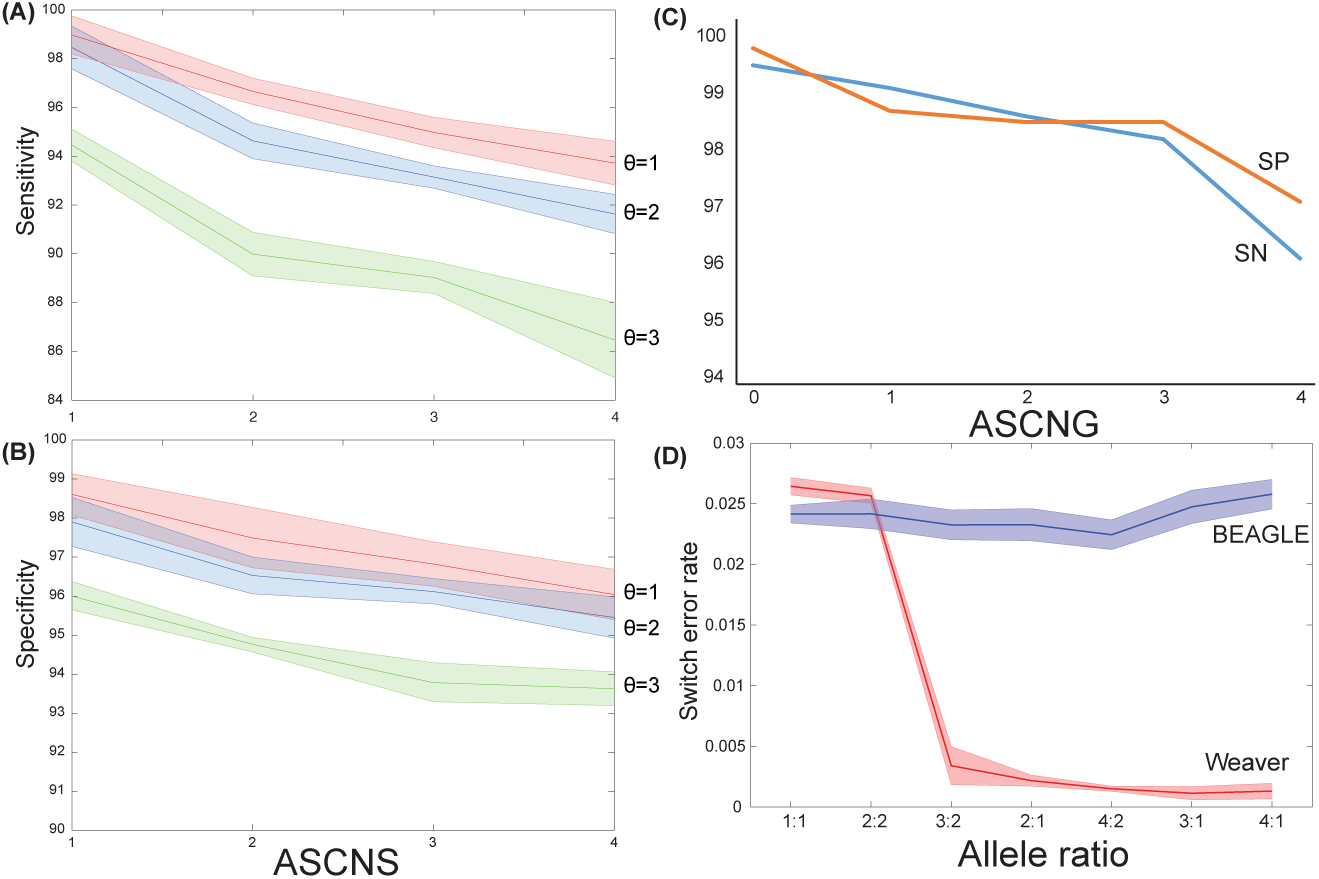
In **(A)** and **(B)**, SN and SP based on results from Weaver are calculated for each ASCNS under different sequencing coverage dispersions (*θ*) using simulation. Random fluctuations are imposed onto initial simulation dataset to create testing dataset with specific dispersion. Both SN and SP decrease with increasing SV CN and dispersion. From observation on real cancer sequencing data, real dispersion is slightly higher than one. **(C)** SN/SP is summarized for each ASCNG from Weaver. **(D)** Switch error rate of Weaver and BEAGLE on simulated datasets with different allele ratios. With imbalanced dataset, error rate of Weaver decreases to less than 0.5%.

#### 3.1.2 Comparison with other ASCNG methods

All CNA methods based on high-throughput technologies including array CGH, SNP arrays and NGS adopt a similar workflow for detection of CNVs and the segmentation is the core part. Signals are used in segmentation, including the signal intensity in array or read counts in NGS, and the b-allele frequency in array or allele frequency in NGS. We first compared Weaver with CNVnator [26] and HMMcopy [27], both designed for partitioning normal genome sequenced by NGS, ignoring allele information. The output of both tools is segmented genome regions with gain, loss or neutral labels, without exact copy numbers. The segmentation results from all three tools were compared to simulation gold standard (if an identified breakpoint is within [-1, 1]kb region of simulated ones, that breakpoint are considered as correct) with both SN (percentage of correctly identified gold standard breakpoints within all reported breakpoints) and SP (percentage of correctly identified gold standard breakpoints within all gold standard breakpoints) calculated. When SV information is omitted (SV ratio = 0), Weaver achieved an average of 80.6% sensitivity and 92.5% specificity in finding copy number change points (Appendix Fig. 2), with increasing SV information, the performance of Weaver gradually improved, suggesting that the advancement of considering CNA together with SV. Even with false SV predictions (SV ratio > 1), Weaver still had accurate results. Also, CNVnator is consistently better than HMMcopy for both SN and SP.

To evaluate the performance on identifying exact ASCNG, we compared Weaver with ASCAT [13] and CNVnator [26]+ThetA [28] (ThetA needs a third party tool to perform segmentation). We converted our sequencing data to logR and BAF values for SNP positions from Illumina HumanOmni2.5 BeadChip (2,015,318 SNP positions genome wide). Overall 43,758 SNPs are within the simulated region. CNVnator+ThetA works on NGS data, but only reports overall copy number. Weaver identified that 97.2% genomic regions have the same copy number with simulation gold standard, while both ASCAT and CNVnator+ThetA had much lower consistency (<20%). These simulation results suggest that it is important to simultaneously consider CNAs and SVs, especially in highly rearranged cancer genomes.

#### 3.1.3 Comparison with other SNP phasing tools

BEAGLE [29] is a statistical phasing method based on a population reference-panel of phased chromosomes. We evaluated Weaver using switching error rate, which is the standard metric for phasing accuracy [30]. The switching error is the proportion of switches in the inferred haplotypes to recover the correct phase in an individual. In our evaluation, we left the phasing information of the testing individual out, and used the rest 1KGP individuals as reference-panel. Overall, Weaver reported an average switching error rate 0.2%, while BEAGLE had an error rate of 3% on regions with imbalanced allele ratio. We have also observed a clear decrease of switching rate for Weaver on dataset with increasing allele imbalance (Fig. 3D). We also attempted to compare with a recently published method HARSH [31]. But HARSH did not finish even on one simulated dataset after more than two weeks, making the comparison difficult to perform.

#### 3.1.4 Simulation from whole chromosomes

We also tested Weaver on simulation dataset derived from whole chr4, chr17 and chr19. Overall, all 52 simulated SVs have been identified by Weaver, with 49 of exact base-pair resolution breakpoint boundaries. The remaining 3 SVs have their breakpoints within low complexity regions and the ‘soft-clip’ strategy (Appendix 5.1.4) failed to identify the detailed breakpoints. However, ‘discordant paired-end’ strategy (Appendix 5.1.4) still identified these 3 SVs with a rough estimation of their breakpoint locations. 100% ASCNS reported by Weaver are consistent with simulation gold-standard.

In terms of relative timing comparing with aneuploidy, all 36 pre-aneuploid SVs in this randomly generated dataset were correctly identified. Weaver labeled 10 SVs as post-aneuploid and 2 of them were incorrect since they were assigned to wrong alleles. These 2 false positive post-aneuploid SVs were actually on the alleles which were not amplified, therefore no timing information would be inferred from them. On ASCNG level, out of 330,988,351bp regions simulated, 2,829,832bp regions (0.85%) had incorrect ASCNG. On the level of overall copy number, ignoring allele information, 1,257,919bp regions (0.38%) had incorrect copy number.

### 3.2 Application to the MCF-7 genome compared with Optical Mapping analysis

We applied Weaver to the whole-genome DNA sequencing data of the MCF-7 breast cancer cell line, with approximately 100X overall coverage and 20X haplotype level coverage. Genome-wide ASCNS and ASCNG are shown in Fig. 4A. 68.3% of MCF-7 genome have imbalanced ASCNG, enabling accurate phasing of SNPs and distal SVs. Weaver identified 546 SVs with 83.3% have copy number greater than 1. Moreover Weaver found 276 post-aneuploid SVs, especially two deletions, *Del*_1_ (chr9:21,837,011-22,081,282) and *Del*_2_ (chr9:21,819,514-21,989,631), within the *MTAP-CDKN2A/B* region (Appendix Fig. 3), where deletions have been frequently observed in various cancers [4, 32]. Weaver found that the short arm of chr9 is triplicated with LOH, having mutually exclusive *Del*_1_ and *Del*_2_, with copy number 2 and 1, respectively. The copy number of deletions implies the region has been amplified twice. Before the first amplification, which changed copy number from one to two, no deletion occurred. After the first amplification and before the second amplification, *Del*_1_ emerged on one chromosome, which was then duplicated in the second round of amplification, leading to two copies of *Del*_1_. Independent from *Del*_1_, *Del*_2_ was formed in approximately the same region of the other chromosome after the first amplification. Such evolutionary information cannot be revealed without SV copy number quantification.

**Figure 4:**
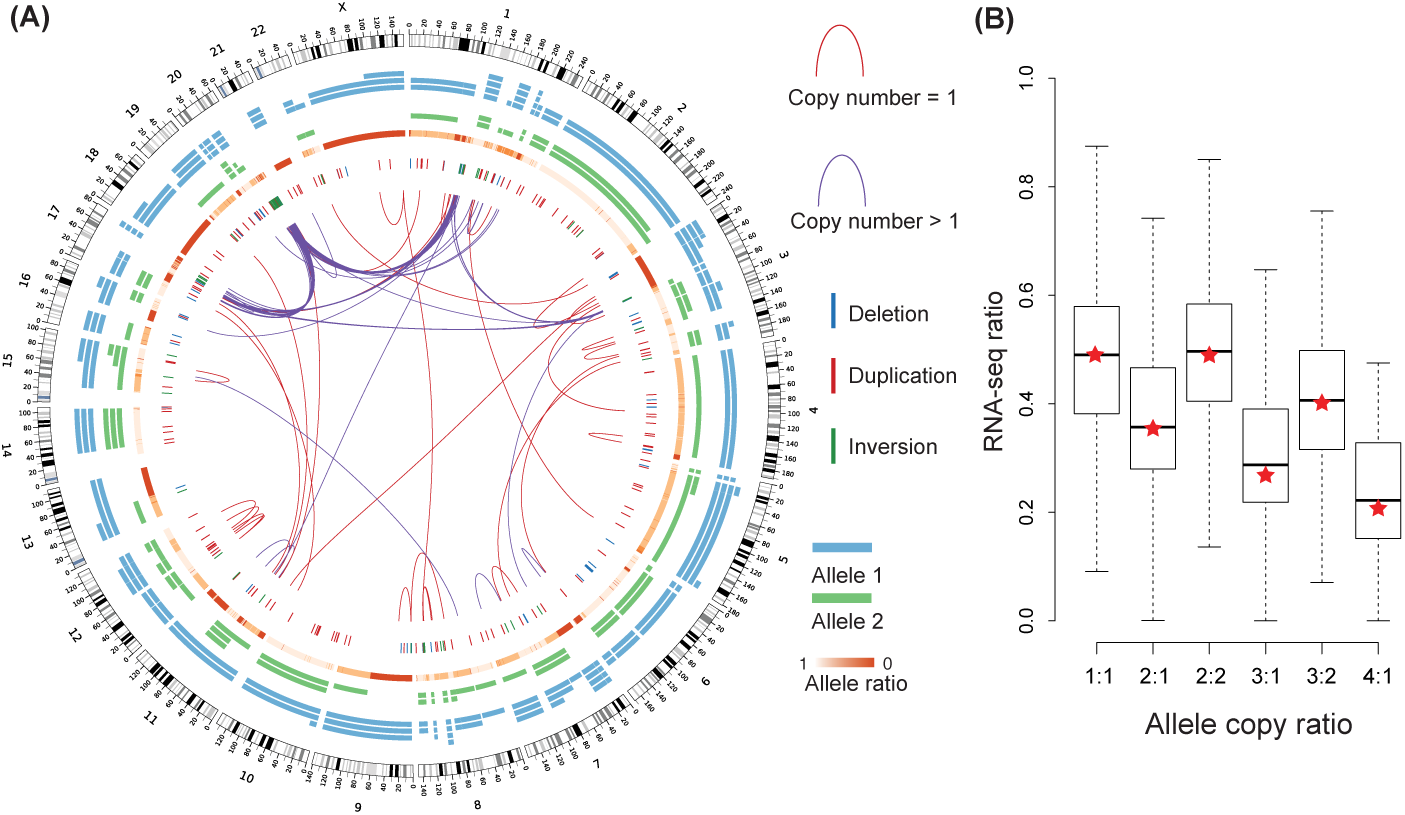
**(A)** Overview of the genomic landscape of MCF-7 cell line. Inter-chromosomal and intra-chromosomal SVs with size larger than 10Mbp are illustrated as red (one copy) and purple (>one copy) lines. Short range deletions, duplications and inversions (imbalanced) are presented as blue, red, and green vertical bars, respectively. chr1, chr3, chr17 and chr20 have interconnected focal amplifications. ASCNGs are plotted as blue and green segments. **(B)** Distribution of allelic expression ratio measured by RNA-seq for different allele copy number categories. Red stars indicate allele copy ratio.

The ASCNG generated by Weaver in MCF-7 demonstrated that detailed cancer genome analysis could help better interpret cancer functional genomic sequencing data. The recent work in HeLa cells has also suggested the need of allele-specific analysis of transcriptome and epigenome data [6]. Fig. 4B shows that ASCNG predicted by Weaver correlates well with the allele-specific expression in MCF-7.

We used Optical Mapping (OM) analysis to compare with the results from Weaver. OM [33–36] is a singlemolecule system that directly constructs large datasets comprising ordered restriction maps (Rmaps; 1 Rmap is 1 mapped DNA molecule) from individual genomic DNA molecules (300kb-2,000kb). After assembly of Rmap datasets into Optical Maps (Rmap contigs), automated detection of structural variants (2kb to multiple megabases) can be discovered spanning large region of genome [37–40], which provides long range linkage information that current NGS approaches are not able to achieve. We selected 268 long range MCF-7 SVs detected by Weaver and built *in silico* ‘cancer reference map’ from these SVs by piecing together 300kbp flanking regions of two breakpoints for each SV. By comparing to the OM result, 235 of the Weaver detected SVs are consistent with OM analysis, suggesting high accuracy of Weaver in identifying SVs. Note that the 33 missed by OM result may not be false positives from Weaver, it is also possible that no Rmaps captured that SV.

An example of OM supporting a tandem duplication (TD) detected by NGS, is shown in Appendix Fig. 4. Two OM Rmaps are shown as blue lines, with red dots represent theoretical cutting cites of restriction enzymes on reference genome and purple dots represent cutting sites that missed by OM. Black number on each segment of OM Rmaps shows the expected length (kb) of OM Rmaps between two cutting sites on the reference and blue number under each segment shows the observed length (kb) of OM Rmaps. As in Appendix Fig. 4, the expected length of OM Rmap covering the TD breakpoint is 23.7kb, while the observed lengths of two OM Rmaps are 22.4kb and 23.1kb, respectively. The strong concordance between expected and observed OM Rmap length on multiple OM Rmaps serves as an independent validation of this TD initially detected by NGS.

### 3.3 Application to the HeLa CCL-2 genome

We applied Weaver to the whole genome sequencing data of the HeLa cells CCL-2 generated by [6]. Haplotype level coverage is approximately 28X. The original study by Adey et al. [6] reported 12 inter-chromosomal SVs, and no large scale intra-chromosomal SV was reported (only deletions and inversions with size < 10kb are reported). However, from our analysis on the same data, we have identified 8 inter-chromosomal and 86 intra-chromosomal SVs (if intra-chromosomal SVs are deletion or tandem duplication type, only those with size > 20kb are reported). Overall, there are 62 genes harboring SV breakpoints.

Genome-wide representation of the Weaver results on HeLa is in Fig. 5A. The large-scale aneuploidy and LOH region is very consistent with [6]. Comparing Weaver’s result with [6], for all genomic regions with copy number profiled, 96.1% have consistent overall copy number estimation between two studies. For ASCNG, the consistency is 97.3% by comparing Weaver output with Table S13 in [6] (and also Fig.1a in [6] for visual inspection of the consistency). Those copy number inconsistent regions include the deletion of *FHIT* gene [41] (Fig. 5B), which is a tumor suppressor gene frequently to have no expression in various human tumors including the HeLa cell line [41–43].

**Figure 5:**
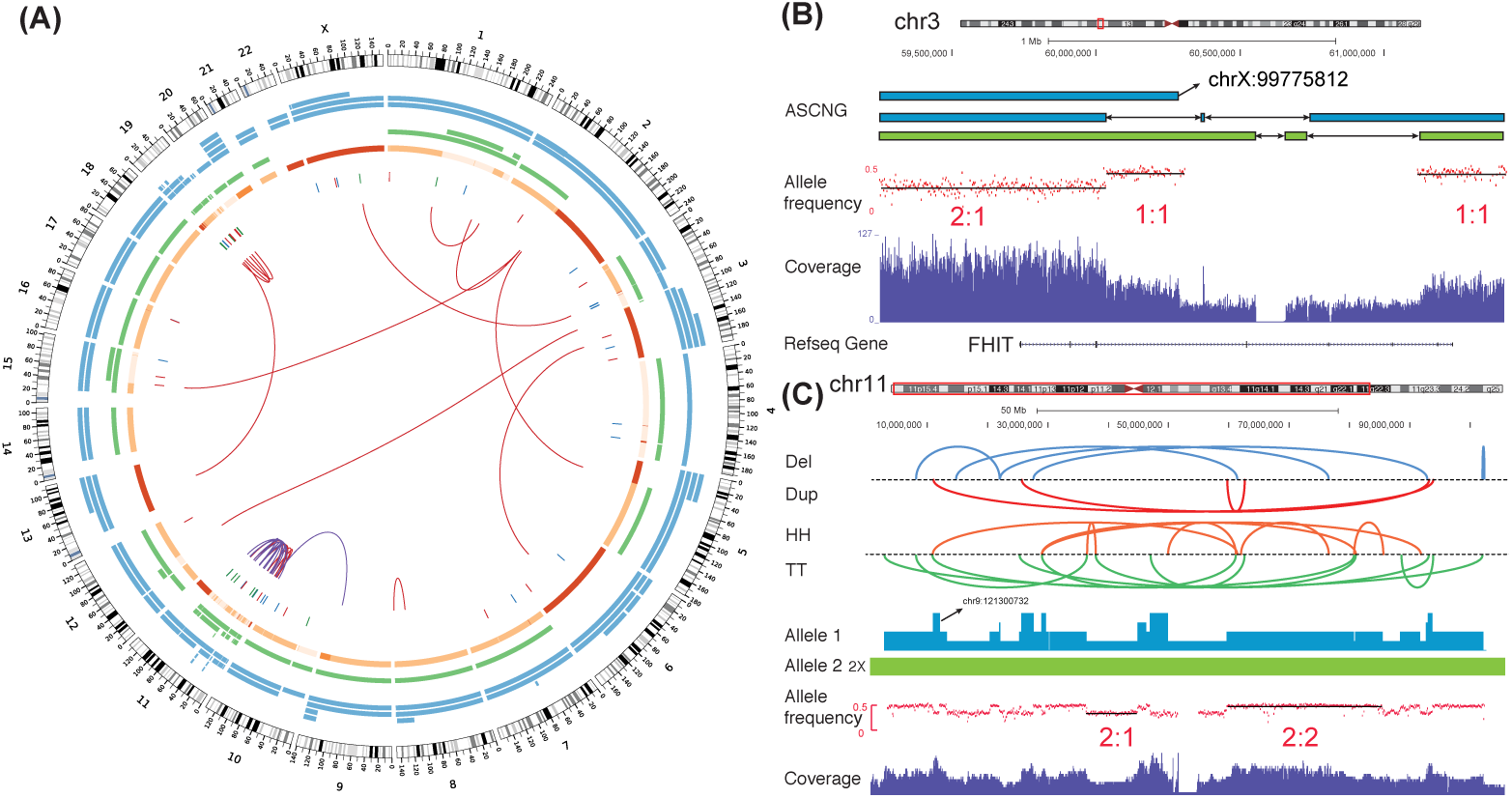
**(A)** Overview of the genomic landscape of HeLa (same legend as in Fig. 4). **(B)** Complex SVs on *FHIT* gene region. Blue/green segments represent chromosomes originated from the same parental allele in normal genome. Allele frequency is plotted by minor allele frequency of germline SNPs. 5 SVs in this small region and all have copy number 1. Adey et al. [6] did not report these SVs or CNAs in *FHIT* gene. **(C)** SVs and ASCN of chr11.

Chr11 and chr19 have undergone extensive amount of SVs (Fig. 5A,C), which have also been reported by [44] in HeLa Kyoto cell line. The SVs are likely to be formed by chromothripsis [45]. With Weaver, we were able to assign different timing for the two chromothripsis events on chr11 and chr19. The chromothripsis on chr11 happened before aneuploidy (since most of the breakpoints have copy number > 1)(Fig. 5C), while chromothripsis on chr19 happened after aneuploidy (since most of the breakpoints have copy number = 1).

### 3.4 Application to 44 TCGA ovarian cancer whole genome sequencing samples

Previous studies have shown that ovarian cancers (OVs) are featured with genetic instability, including recurrent nonrandom chromosomal abnormalities, multiple chromosomal losses and gains, and the presence of marker chromosomes [46–49]. TCGA provided a detailed catalogue of genomic aberrations in OV [50], suggesting that the degree of somatic CNAs in OV is strikingly high comparing with other tumors. We applied Weaver on 44 high coverage (>15X on haplotype level) TCGA OV samples. We found that these samples typically do not have complex subclones, suggesting the feasibility of applying Weaver. The overall rearrangement maps for all the samples is in Appendix Fig. 5.

Genome-wide representation of Weaver result on one sample, TCGA-36-1571, is in Appendix Fig. 6A. There are two groups of highly inter-connected chromosomes: chr4-chr22 and chr6-chr14 (Appendix Fig. 6B-C). By calculating the detailed copy number of involved SVs and genomic regions, the chr4-chr22 group showed signatures of chromothripsis of multiple chromosomes, while chr6-chr14 group showed extensive focal copy number gains and is most likely to be formed by progressive process other than a single catastrophic chromosome shattering event. Indeed, chr6-chr14 region has high number of fold-back inversions, which are fundamental to progressive rearrangements driven by breakage-fusion-bridge (BFB) repair [17, 51, 52]. Interestingly, *FOXG1* gene is proximal to the BFB fold-back inversion site on chr14 and highly amplified (Appendix Fig. 6C). It was reported that the over-expression of *FOXG1* contributes to TGF-*β* resistance in OV, leading to loss of growth inhibitory response to TGF-*β*, a common feature of epithelial cancers [53].

**Figure 6:**
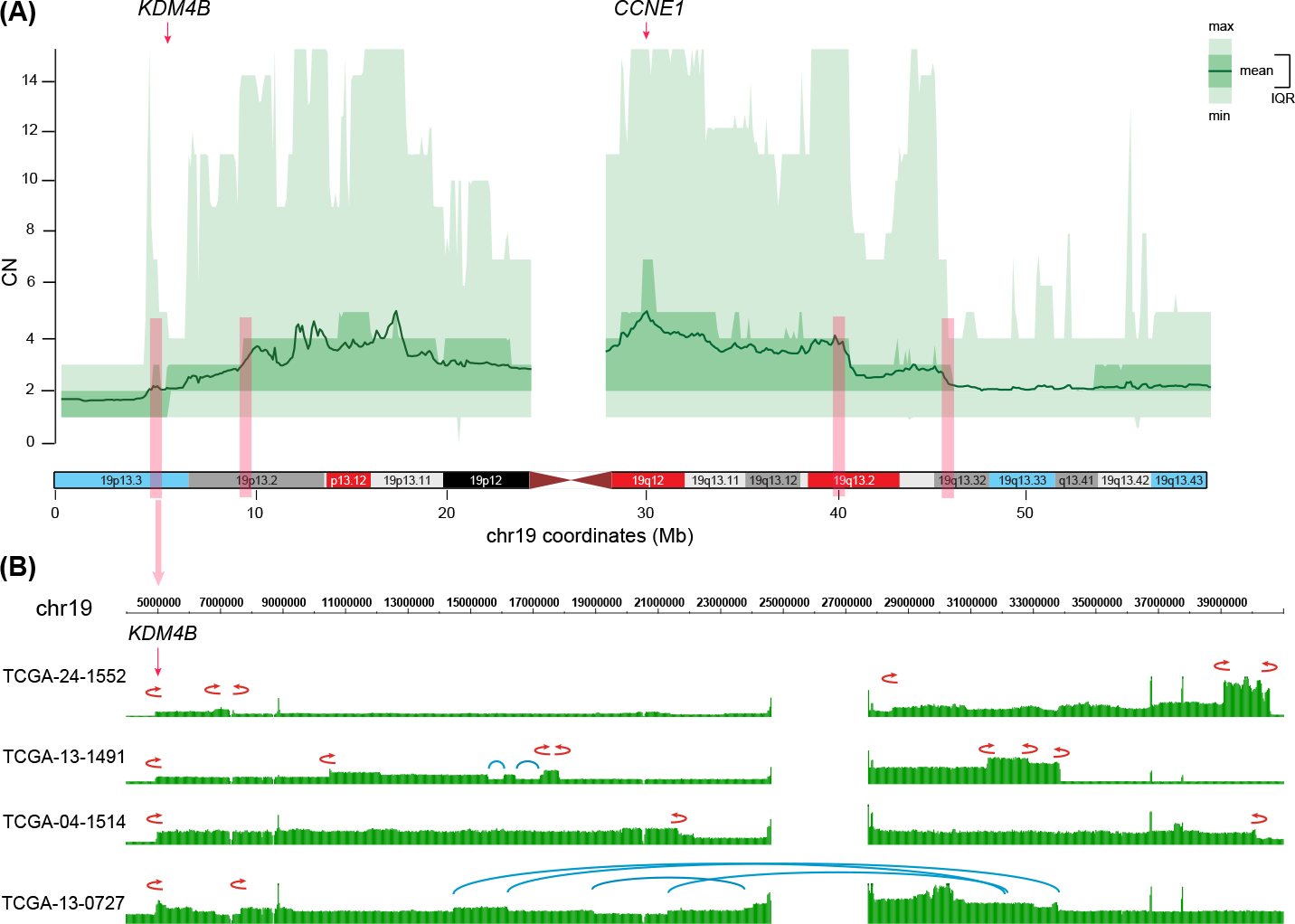
Copy-number landscape of chr19 is altered by recurrent SV. **(A)** The overall copy number profile across 44 OV samples. The chromosome bands are highlighted with red and blue, representing the frequently amplified and deleted regions from GISTIC analysis of TCGA OV array data [50]. *CCNE1* gene is within most significantly amplified region. Four pink tracks represent regions on which gross copy number profile has changed and all four regions have recurrent FBIs. **(B)** Four OV samples with recurrent FBI at *KDM4B* are illustrated. Red fold-back arrows represent FBIs while blue lines represent other intra-chromosomal SVs.

Fig. 6A shows the overall copy number profile of chr19 across 44 OV samples. Gene *CCNE1* is within the most significantly amplified region. Amplification of *CCNE1*, which encodes cyclin E1, is associated with primary treatment failure in women with OV. *CCNE1* copy number is validated as a dominant marker of patient outcome in OV [54]. Previous studies have reported that *CCNE1* amplification is one of the most common focal copy number change events in OV [50, 55].

One small region on 19p13.3 (4.6M-6.7M) is enriched with fold-back inversions that lead to amplification of 19p13.2. Especially TCGA-04-1514, TCGA-24-1552, TCGA-13-1491 and TCGA-13-0727 (4 out of 44 deep sequencing OV samples analyzed) have FBIs with breakpoints within a < 60kbp region, as shown in Fig. 6B. Interestingly, the breakpoints are right around *KDM4B* from *KDM4* protein family which are demethylases that target histone H3 on lysines 9 and 36 and histone H1.4 on lysine 26 [56]. Various studies have shown that *KDM* proteins, including *KDM4B*, are over-expressed in different types of tumors and are required for efficient cancer cell growth.

Taken together, our results demonstrated the potential of Weaver to refine the analysis SVs and CNAs in complex cancer genomes using whole-genome sequencing data.

## 4 Discussion

Genomes of somatic cells undergo dramatic and complex genetic changes during cancer development, including point mutations, SVs, large-scale gain or loss, and even aneuploidy. Genome aneuploidy is a common feature of cancer cells. Recent pan-cancer analysis based on whole-genome sequencing data estimated that over one third of the tumors have whole-genome duplication, and the proportion can reach over 60% in some types of cancer [4]. In addition, genome aberrations caused by SVs and copy number changes are a common feature of a wide variety of neoplastic lesions. Recent advances in NGS technologies have provided us with an unprecedented opportunity to better characterize these different genomic changes in cancer. However, even though methods have been separately developed to identify SVs and CNVs using NGS reads [5, 6, 9, 11, 57–59], no algorithm is currently available to simultaneously study SVs and CNVs in aneuploid cancer genomes. Therefore, Weaver represents the first method that quantifies allele-specific copy numbers of SVs in cancer genomes and provides a more integrative solution to study complex cancer genomic alterations.

We expect that Weaver will be very useful to refine the analysis of the existing datasets in large-scale projects such as TCGA and ICGC, which were mostly sequenced using short read NGS technology. The algorithm in Weaver is not restricted to short reads and can adapt to data from longer read sequencing technology. However, it will remain difficult to completely elucidate those very large complex SVs in cancer genomes, especially when the breakpoint regions caused by SVs contain highly repetitive sequences.

There are a number of areas that the Weaver algorithm can be further improved. Copy number neutral events such as balanced inversions are not currently handled in Weaver. Although Weaver sets no limit on maximum CN, its accuracy in quantification SVs is naturally hampered in highly repetitive regions, either in the reference genome or in the cancer genome. In the case of MCF-7 genome, chromosomes 3, 17 and 20 have regions with higher than 100X copies, and the estimation on CN and phasing of SVs within those regions may be less reliable. In addition, even though the probabilistic graphical model employed in Weaver is generic to consider complex tumor subclones caused by intra-tumor heterogeneity, current version of Weaver only works for samples with a dominating tumor cell population (which can be estimated by tools such as ABSOLUTE) with possible normal cell contamination. However, recently a number of new algorithms have been developed to specifically identify subclonal structure of tumor cell populations [28, 60–65]. Method like TITAN [66] was also developed to estimate ASCN alterations in the a mixture of tumor cell population, although TITAN does not handle complex SVs. Nevertheless, the results from these algorithms that identify tumor subclone architecture are complementary to what Weaver can achieve. However, new methods are needed to quantify ASCN of SVs and understand how complex SVs interact in the context of a mixture of aneuploid tumor cell population to reconstruct the evolutionary history of tumor genomes.

